# Cracking the code of co-authorship networks geo-temporally using interpretable machine learning

**DOI:** 10.1101/2025.03.05.641725

**Authors:** Swapnil Keshari, Zarifeh Heidari Rarani, Akash Kishore, Jishnu Das

**Affiliations:** Department of Computational Biology, University of Pittsburgh, Pittsburgh, PA, USA; Department of Immunology, University of Pittsburgh, Pittsburgh, PA, USA; Center for Systems Immunology, University of Pittsburgh, Pittsburgh, PA, USA; The Joint CMU-Pitt Ph.D. Program in Computational Biology, Carnegie Mellon University and University of Pittsburgh, Pittsburgh, PA, USA

## Abstract

An exponential growth in the scientific literature necessitates the development of highly scalable computational tools that can effectively analyze and distill insights from complex, interconnected research landscapes. We introduce Distributed, Interpretable, and Scalable computing for Co-authorship Networks (DISCo-Net), a robust and scalable tool engineered to curate and examine large-scale co-authorship networks by harnessing the power of distributed computing and advanced relational database queries. We use DISCo-Net to analyze co-authorship networks derived from millions of papers in the life sciences and physical sciences over more than two decades. Using a range of deep learning approaches, we surprisingly found that pre-trained zero-shot embeddings from a sentence transformer better captured global co-authorship relationships than a complex graphical attention transformer. Even more surprisingly, a simple interpretable Term Frequency-Inverse Document Frequency (TF-IDF) model performed as well as the Bidirectional Encoder Representations from Transformers (BERT) model. Through topic modeling on TF-IDF document descriptors, we identified nine major research areas prevalent globally over the past 24 years and captured topic-specific shifting trends in scientific output. Our study draws an innovative parallel between collaborative research networks and genomic regulatory structures, applying genomics data analysis methodologies to uncover patterns in global scientific collaboration. This approach reveals interpretable alignments between research interests and human developmental stages, while also identifying emerging influential players in the global research landscape. The findings highlight potential far-reaching consequences of current funding challenges, particularly in the U.S., and offer actionable insights for optimizing resource allocation and fostering innovation in an interconnected global scientific community.

## Introduction

Humans naturally connect through shared interests and expertise, often spanning diverse fields in this interdisciplinary world. The practice of sharing formal research, central to scientific progress, dates back to 1665 with the founding of *Le Journal des Sçavans*^1^ and *Philosophical Transactions*^2, 3^, the first journals aimed at keeping scientists informed of each other’s work. While most early scientific articles reflected solo efforts, Beaver and Rosen^4^ demonstrated that collaborative science originated as early as the 17^th^ century. They introduced a “comprehensive framework” to study collaboration in scientific articles, highlighting co-authorship, and challenging the prevailing belief that scientists worked in highly isolated communities and subcommunities.

A deeper understanding of human relationships offers valuable insights into societal conduct, including scientific ones. In a first-of-its-kind study in 1933, Monero^5^ introduced sociograms to analyze social interactions leveraging a network-based approach. He emphasized “the need for determining the nature of fundamental structures forming the networks to solve problems caused by group attraction, repulsion or indifference”. Over subsequent decades, significant progress transpired in probabilistic network theory, random graph models^6^, and the discovery of scale-freeness in real-world networks^7^, solidifying “network science” as a major field of endeavor. Such advances in network science coupled with derived insights from them into human behavior have profoundly shaped scientific social contracts. These policies address critical questions such as: Who conducts research? Who funds it? Who benefits from it? How does attention to different research topics evolve over time and geographical locations?

Established in 1848, the American Association for the Advancement of Science (AAAS)^8^ pioneered formal scientific outreach policies and was open to all. This was followed by the founding of the National Academy of Sciences in 1863^9^, backed by Bache and his exclusive group. They advocated for federal funding of scientific research and sought to restrict scientific practice to those conducting “high-quality research”. With time such decision making has evolved, increasingly driven by research itself rather than individual or group opinions and ideas. Analyzing scientific relationships from a network perspective provides valuable insights, enabling policymakers to make informed decisions that create a positive feedback loop, accelerating scientific progress and reshaping policies. Today, U.S. federal agencies like the NIH and NSF prioritize collaborative research^10^, drawing on diverse intellectual resources. This approach not only provides new researchers with greater visibility and access to resources but also mitigates the concentration of power and prevents the stratification of individuals in shaping scientific fields.

Co-authorship networks have been extensively studied for insights into scientific collaboration, with traditional analyses focusing on topological features. However, these analyses have primarily focused on highly cited papers and individuals within specific research fields and regions. Moreover, the lack of well-organized data coupled with the increasing scale and complexity of these networks constrained these studies by geography, time, or discipline. These limitations have restricted the ability to capture large scale relationships across the network. Combined with rising computational and memory demands, they have created a significant gap in fully leveraging the potential of network analysis methods.

Addressing these gaps, we provide an easy-to-use python package “**D**istributed, **I**nterpretable and **S**calable computing for **Co-**authorship **Net**work” framework DISCo-Net. DISCo-net addresses a long-standing need to incorporate processing, modeling and inference for networks. DISCo-Net can readily incorporate other ML models and datasets, providing a one-stop framework for network science driven policy making. Using DISCo-Net, this study performs a comprehensive analysis of scientific collaborations using a novel network-based framework. We adapted extremely scalable and efficient data processing algorithms to fetch and process over 2 decades of scholarly data in the life sciences and physical sciences to construct large-scale co-authorship networks. Following this, we use traditional networks-based approaches to analyze the nature and patterns of co-authorship. As expected, co-authorship networks over time grew and maintained their scale-free properties. But beyond providing these expected findings, they did not provide novel insights. Hence, we switched to modern deep learning methods, specifically graphical attention networks^11^ (GATs). To enable prediction of collaborations within co-authorship networks, we tasked the GATs with link prediction problem. Link prediction plays a foundational role in our ability to assess the training, utility, and effectiveness of graphical models. Next, this study delves deeper into dissecting the superiority of GATs and characterize the role of low dimensional embeddings (BERT^12^) in GATs success across time, communities and sub-communities. Since interpretable inferences are paramount for policy level inputs, we then use Term Frequency-Inverse Document Frequency (TF-IDF) to infer global scientific research across time. Topic modeling^13^ of TF-IDF-derived terms reveals temporal global scientific research trends. Overall, we find nine major research areas prevalent globally over the past 24 years and capture shifting trends in scientific output.

Our analysis reveals a striking parallel between the structure of collaborative relationships in research and the architecture of genomic regulatory networks. By applying methodologies from genomics data analysis to this context, we first uncover interpretable patterns that illuminate the alignment of research interests with various levels of human development. This approach not only highlights global research gaps but also yields actionable insights for the strategic allocation of research expenditures. The synergy between these seemingly disparate fields—collaborative networks and genomics—provides a novel lens through which to view and optimize the landscape of scientific collaboration and resource distribution. Building on this, we further leverage analytical methods from genomics to identify emerging players in global collaborations that exert asymmetric influence on specific research topics. This approach highlights the potential far-reaching consequences of current funding challenges, particularly in the U.S., which may impact not just U.S.-led biomedical research but the global scientific community. Our findings have significant implications for science policy and funding strategies in an interconnected and rapidly changing global landscape, offering valuable insights for policymakers and research institutions to optimize resource allocation and foster innovation.

## Results

### Comprehensive analysis of scientific collaborations across time and domains reveals shifts in collaboration patterns and research priorities

Over the past two decades, life sciences research has expanded exponentially, producing nearly two million publications and fostering unprecedented international collaboration. This growth underscores the necessity of protecting and strengthening research networks, not only in the U.S. but worldwide. Data-driven disciplines, spanning computer science, natural language processing (NLP), and computational biology, serve as the backbone of modern science, enabling researchers to extract meaningful insights from vast datasets and predict emerging trends. By analyzing yearly co-authorship networks—where nodes represent authors and edges signify collaborations— using interdisciplinary science, we uncover the evolving structure of scientific communities (Figs. 1A-B, Supplementary Figs. 1A-B). Analyzing these networks reveals shifts in global research priorities, patterns of collaboration, and the resilience of scientific endeavors against political and economic pressures. We first investigate the extent to which local contextual information alone captures global patterns of scientific collaboration over time, by inferring semantic information from publication titles using advanced NLP techniques. While modeling interactions between authors across communities effectively captures both local and global relational information, we conduct a comparative assessment of models that leverages integration of title-driven embeddings and local network metrics to determine how textual data enhances structural features in predicting collaborations. This analysis suggests that simpler models not only outperform others in preserving semantic integrity and interpretability but also offer valuable insights into thematic shifts in scientific inquiry through topic modeling. Additionally, temporal analysis of these interpretable topic models reveals significant disparities in the pace of scientific progress over time, with further implications yet to be fully explored.

**Figure 1.**
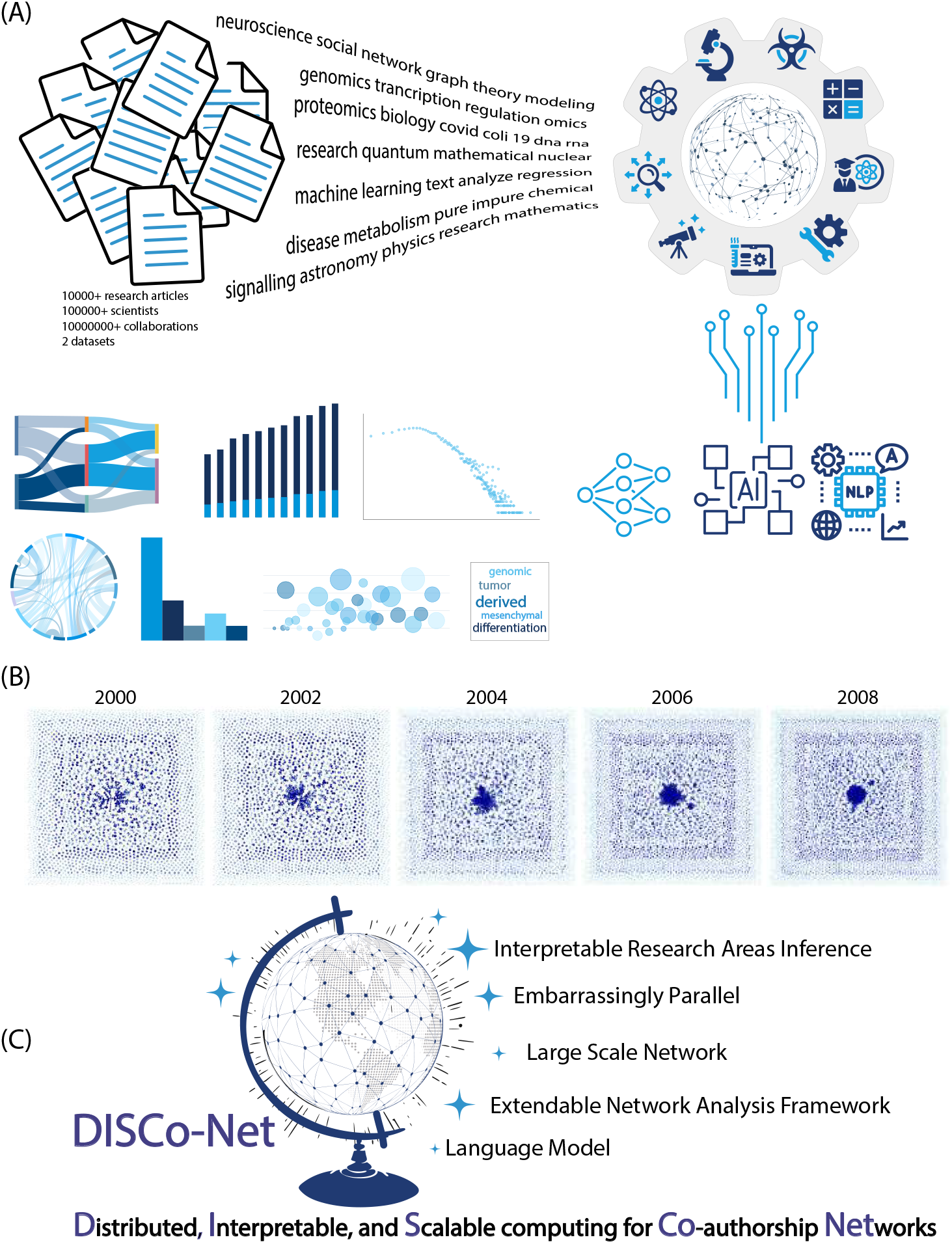
Co-authorship networks in the physical and life sciences over 2 decades. A – Overview of the study B – Structure and topology of co-authorship Networks over the years 2000, 2002, 2004, 2006, and 2008 C – Structure of DISCo-Net – Distributed, Interpretable, and Scalable computing for Co-authorship Networks, a tool for the comprehensive analyses of global scientific collaborations across time and domains.

Then, borrowing methodologies from genomic data analysis equipped us for clustering the countries based on their research outputs across topics and time to shed light on how research priorities are linked to a country’s development status. Notably, our method also provides a way to approximate Human Development Index (HDI) ranks for regions lacking official data by analyzing their research repertoires. It also helps identifying the research footprint of every single country. This provides valuable insights for developing tailored scientific policies and gaining a deeper understanding of global research trends. Analyzing collaboration patterns through both ‘within-country’ and ‘cross-country’ partnerships, we demonstrate how international cooperation trends evolve over time, despite the prevalence of nation-first policies. We capture nuanced temporal trends, highlighting the shifting roles of traditional research leaders and emerging hubs. The scale and interdisciplinary nature of scientific data demand robust, scalable computational frameworks. Currently, no such framework exists for analyzing the dynamics of global scientific collaboration. To address this, we developed DISCo-Net, a distributed computing package that streamlines data retrieval, processing, and analysis. It operates in four key steps: (1) Fetching and preprocessing relevant data, (2) Constructing co-authorship networks and constructing features to represent authors, (3) Training graph-based algorithms for prediction, and (4) Extracting interpretable insights (Fig. 1C). By providing a comprehensive view of the global research landscape, DISCo-Net reveals how collaboration patterns, research priorities, and economic factors shape scientific progress, offering actionable insights for research policy and equitable international cooperation.

### Unravelling relational patterns and collaboration structures using deep learning

We first investigated co-authorship networks across different years using topological network metrics. These include network diameter, average node betweenness, average degree centrality, and global clustering coefficient (Supplementary Fig. 2A). Network diameter measures the largest distance between any two nodes in the network, providing insights into the overall connectivity and extent of co-authorship collaboration. Average node betweenness identifies influential nodes that contribute significantly to the network’s overall connectivity and resilience. We also analyze average degree centrality to understand the typical number of connections each author has, highlighting the overall level of direct collaboration within the network. The global clustering coefficient of a node, quantified as the proportion of connections among its neighbors, is also determined. This coefficient quantifies the extent to which the neighbors of a particular node are interconnected with each other. The average of this value across all nodes serves as a reliable indicator for detecting the existence of community structures within a network.

While the sizes of these networks increased monotonically over time, in line with expectation, these network metrics remained largely constant over time (Supplementary Figs. 2B-C). Despite their utility in characterizing network properties, they do not capture the dynamic aspects of link formation and the evolution of collaborative efforts over time. Notably, we observe a consistent scale-free structure in co-authorship networks in both the life and physical sciences (Supplementary Figs. 2 B-C), consistent with prior expectation. Over time, these networks exhibit remarkable stability in their network topological and scale-free properties. The persistence of high-degree hubs highlights their critical role in maintaining network connectivity and robustness. These findings necessitate a move away from conventional topological frameworks and further explore predictive models to uncover hidden patterns and develop robust predictors for link formation and collaborative efforts.

Even though co-authorship networks are scale free, there is still diversity in author groups and their properties, which topological analyses do not capture. We use a stratification approach based on network topological properties to stratify this. Specifically, we subset authors (nodes in the network) in four groups pivoting on either their degree and published paper counts or degree and node betweenness (Figs. 2A-B). While degree captures connectivity of an author, paper counts quantify productivity of the author and node betweenness reflects the nature or quality of collaborations. Together, these 2 metrics define 4 groups corresponding to different authorship patterns based on productivity and connectivity. For example, group 3 in papers published vs degree plot consist of authors who have very high connections (i.e. degree) and very high publications. Another example can be people in group 2 when looking at the node betweenness vs. degree plot. While such scholars are highly connected, they aren’t as central to the research network, representing intramodular hubs. Another such real-world instance which this stratification captures is nodes with high betweenness, and low degree centrality formed critical bridges in the network, often linking otherwise disconnected clusters, while nodes with low centrality measures represented more specialized, isolated collaborations (Figs. 2A-B).

**Figure 2.**
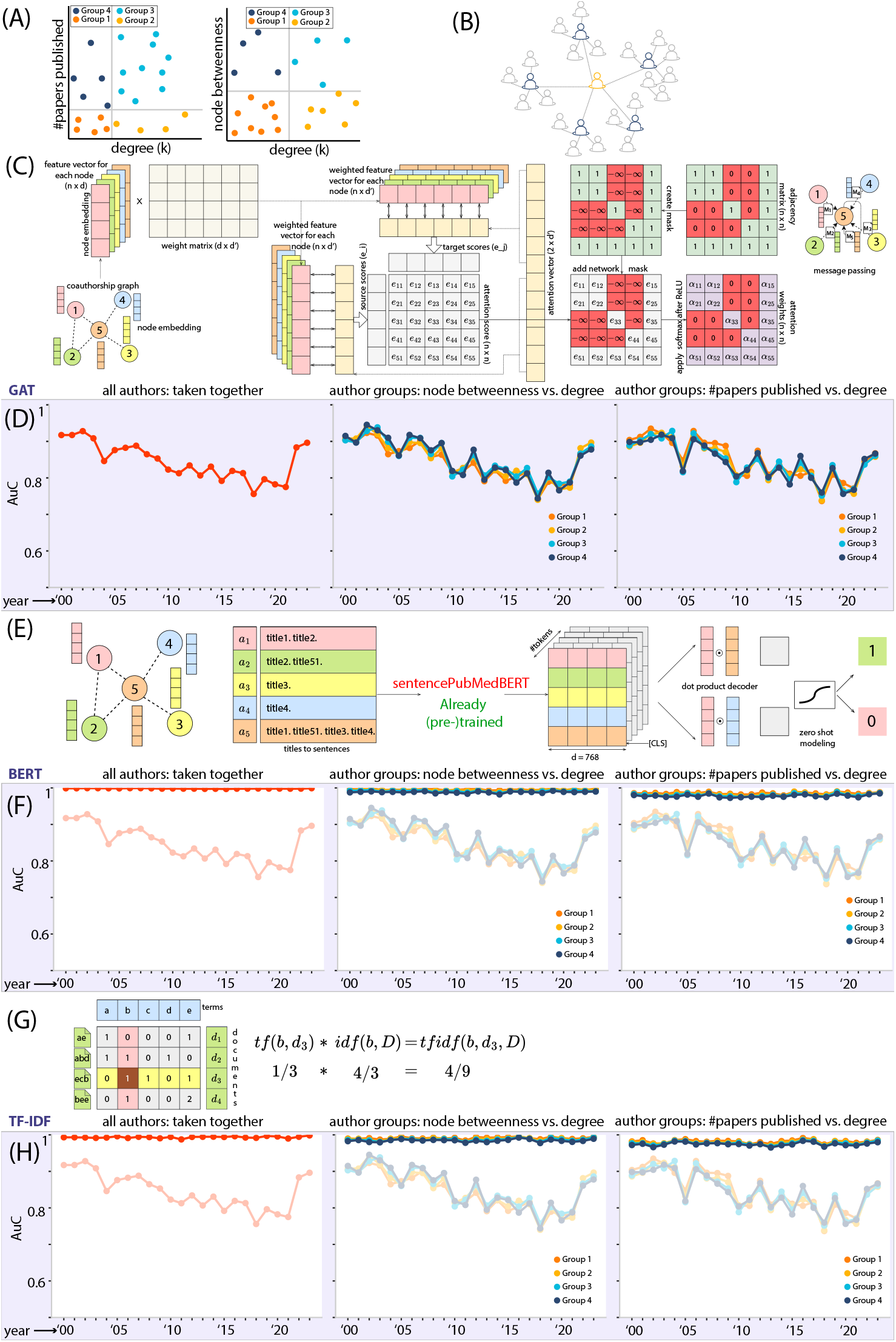
Exploring co-authorship patterns over time and domains across graphical attention transformers, Title-driven embeddings and local information. A – Defining author (nodes in the network) subsets into four groups based on either their degree and published paper counts or degree and node betweenness. B – Architecture of node degree, and node betweenness. C – Architecture and structural details of the GAT. D – AUCs of co-authorship link prediction by the GAT model from 2000-2023 in the life sciences. Panels reflect performance overall, performance for different co-authorship patterns as defined by stratification using degree and node betweenness, performance for different co-authorship patterns as defined by stratification using degree and number of papers published. E – Architecture and structure of the BERT model. F – AUCs of co-authorship link predictions by the BERT model from 2000-2023 in the life sciences. Panels reflect performance overall, performance for different co-authorship patterns as defined by stratification using degree and node betweenness, performance for different co-authorship patterns as defined by stratification using degree and number of papers published. G – Architecture and structure of the TF-IDF model H – AUCs of co-authorship link predictions by the TF-IDF model from 2000-2023 in the life sciences. Panels reflect performance overall, performance for different co-authorship patterns as defined by stratification using degree and node betweenness, performance for different co-authorship patterns as defined by stratification using degree and number of papers published.

Given the complexity and intricacy of the co-authorship structures, we explored GATs to analyze co-authorship networks, especially for link prediction tasks. GATs are particularly well-suited to capture relational information among entities by enabling more intelligent message passing where the convolution process assigns greater importance to certain neighbors over others. The motivation behind coupling GAT with a graphical autoencoder stems from the need to enhance the model’s ability to capture both global and local structures within the network. This architecture allows for effective message passing by integrating the advantages of attention mechanisms, which prioritize significant relationships, with autoencoder-based dimensionality reduction to uncover latent features within the network (Fig. 2C). Thus, GATs capture interactions between authors across communities both based on collaborative patterns (quality) and productivity (quantity of publications). Across collaboration patterns and productivity, GATs are reasonably accurate at capturing co-authorship patterns between author groups across the life sciences and physical sciences (Fig. 2D, Supplementary Fig. 2D, AUC ∼ 0.75). This underscores that GATs are not only capturing global signal, but they have knowledge about local connectivity structure too, something where conventional investigation fails drastically.

Additionally, GATs model performance remains consistent across years. Given that GATs leverage a balance of global and local information from network nodes for prediction, we hypothesized that the predictive signal comes from a balance of the pre-trained embeddings and the structural complexity introduced by GATs. We next chose to conduct methodical experiments to disentangle the contributions of these 2 components and assess their impact on the overall accuracy of link prediction within co-authorship networks.

### Rethinking network modeling: when structure becomes secondary

Building upon the insights gained from deep learning-based methods, we challenged the traditional assumption that topology is indispensable for effective network modeling. Specifically, we tested whether a co-authorship network could be accurately modeled without considering its underlying topology—effectively treating the problem as a zero-shot network modeling task. We harnessed the power of sentence-transformers, a variant of Bidirectional Encoder Representations from Transformers (BERT), specifically tailored to generate high-quality sentence embeddings and enhancing semantic understanding in biomedical natural language processing (NLP) tasks. BERT is a pretrained language representation model developed by Google that uses a transformer-based architecture to process text in a way that considers the context of each word in both directions (left-to-right and right-to-left) in all layers. While BERT can perform sentence-pair regression, it does not provide any independent sentence embeddings, making it computationally intractable to compare sentences (*O*(*n*^2^)) in a large dataset. Hence sentence transformers, leveraging siamese network architecture, are generally used to generate fixed-size embeddings for input sentences. Semantic similarity between sentences can then be found by utilizing similarity measures like cosine similarity or Manhattan / Euclidean distance.

We used a zero-shot model, a technique that allows the model to make predictions on previously unseen data without task-specific training. Our objective was to determine whether the complexity of the GAT architecture was necessary or if a simpler formulation could achieve comparable results. Here, we generated embeddings for each author using Sentence BERT language model (Fig. 2E). In contrast to BERT, sentence BERT captures the context of the entire sentence and can generate corresponding embeddings. We used a model that was pre-trained on Pubmed articles and fine-tuned on MS-MARCO, to ensure that it has context specificity. We encoded paper titles for each author in data via Sentence BERT to create dense vector representations and use [*CLS*] token which captures whole sentence context. On comparing GAT-based predictions with those derived solely from pre-trained embeddings, we discovered that the structural complexity introduced by GATs did not necessarily improve predictive accuracy. Surprisingly, we found that high-quality embeddings alone were sufficient to achieve robust link prediction, as evidenced by the *AUC* ≈ 1 across both the life and physical sciences domains across over 2 decades (Figs. 2F and Supplementary Fig. 2E). Our findings suggest that a well-structured zero-shot model combined with effective embeddings significantly outperform more complex network structures incorporating attention. Our analyses highlight the effectiveness of sentence transformers with meaningful embeddings in achieving almost perfect link prediction. The performance remains constant across the 4 groups of authors defined earlier based on connectivity and productivity in both the life sciences and physical sciences (Figs. 2F and Supplementary Fig. 2E), highlighting that the embeddings capture meaningful information across different co-authorship patterns. Our results present a major conceptual advance demonstrating that just local contextual embeddings reflected in titles are good enough to understand global co-authorship patterns across domains (life and physical sciences), over two decades of scientific research (2000-2023), spanning different connectivity patterns (4 quadrants defined based on productivity and network centrality).

### An interpretable model is as informative of co-authorship patterns as BERT embeddings

While BERT-based encodings work extremely well, a key limitation of the approach is their inability to provide meaningful interpretability. Thus, we implemented Term Frequency – Inverse Document Frequency (TF-IDF) for generating document embeddings (Fig. 2G). TF-IDF treats each paper title as a document and considers all titles published within a year as the document corpus. Unlike BERT-based embeddings, TF-IDF does not capture the semantic nuances of sentences but offers a distinct advantage in terms of interpretability. Each dimension of a TF-IDF embedding corresponds to a specific term, providing reasonable dimensions, with its importance evaluated relative to the entire corpus. Our analysis demonstrated that TF-IDF still performed as well as the Sentence BERT model with complex embeddings across time (Figs. 2H, Supplementary Fig. 2F). This indicates that while high-quality embeddings are effective, TF-IDF is as informative of co-authorship patterns across domains (life and physical sciences), over two decades of scientific research (2000-2023), spanning different connectivity patterns (4 quadrants defined based on productivity and network centrality). The interpretability of TF-IDF embeddings allows for a clear understanding of the terms influencing predictions, making it particularly useful for qualitative analyses. This capability is evident in our analysis of research trends across different countries over time, something we dissect more carefully in the following section. By analyzing both pre-trained embeddings and TF-IDF-generated embeddings, our study reveals that while sophisticated models like GATs offer certain advantages, their complexity is not always necessary. Effective embeddings, whether generated by advanced models like BERT or simpler methods like TF-IDF, can provide the necessary structure for accurate predictions, making them a versatile tool in the analysis of co-authorship networks.

### Topic modeling of TF-IDF-derived terms reveals temporal global scientific research trends

Extracting immediate interpretable insights summarizing this huge research landscape is challenging. We aimed to identify key biomedical research areas and analyze how their output has evolved temporally, forming foundation to aid data-driven scientific policies. To derive principal research areas from the TF-IDF analyses, we first embed these words from all 24 years and clustered embeddings to uncover semantically similar word clusters. Next, applying c-TF-IDF we identify high frequency important words (or terms) distinguishing each cluster (or research areas) with rest all others. Strikingly, top words in each cluster consistently reflected a meaningful theme, enabling interpretable cluster annotation to define principal research topics across 24 years (Fig. 3A, Supplementary Fig. 3A). Specifically, we identified nine major focus topics, including “Viral Infections & Public Health” (Topic 3), “Climate Change & Environmental Impact” (Topic 0), “Chromatin Structure & Genomics” (Topic 7), “Metabolism, Diabetes & Disease Risk” (Topic 4), “Neuroscience, Cognition & Brain Function” (Topic 1), and “Quantum Imaging & Optical Physics” (Topic 2), among others (Supplementary Table 1). Similarly, for the physical sciences dataset, we identified meaningful topics (Supplementary Table 2).

**Figure 3.**
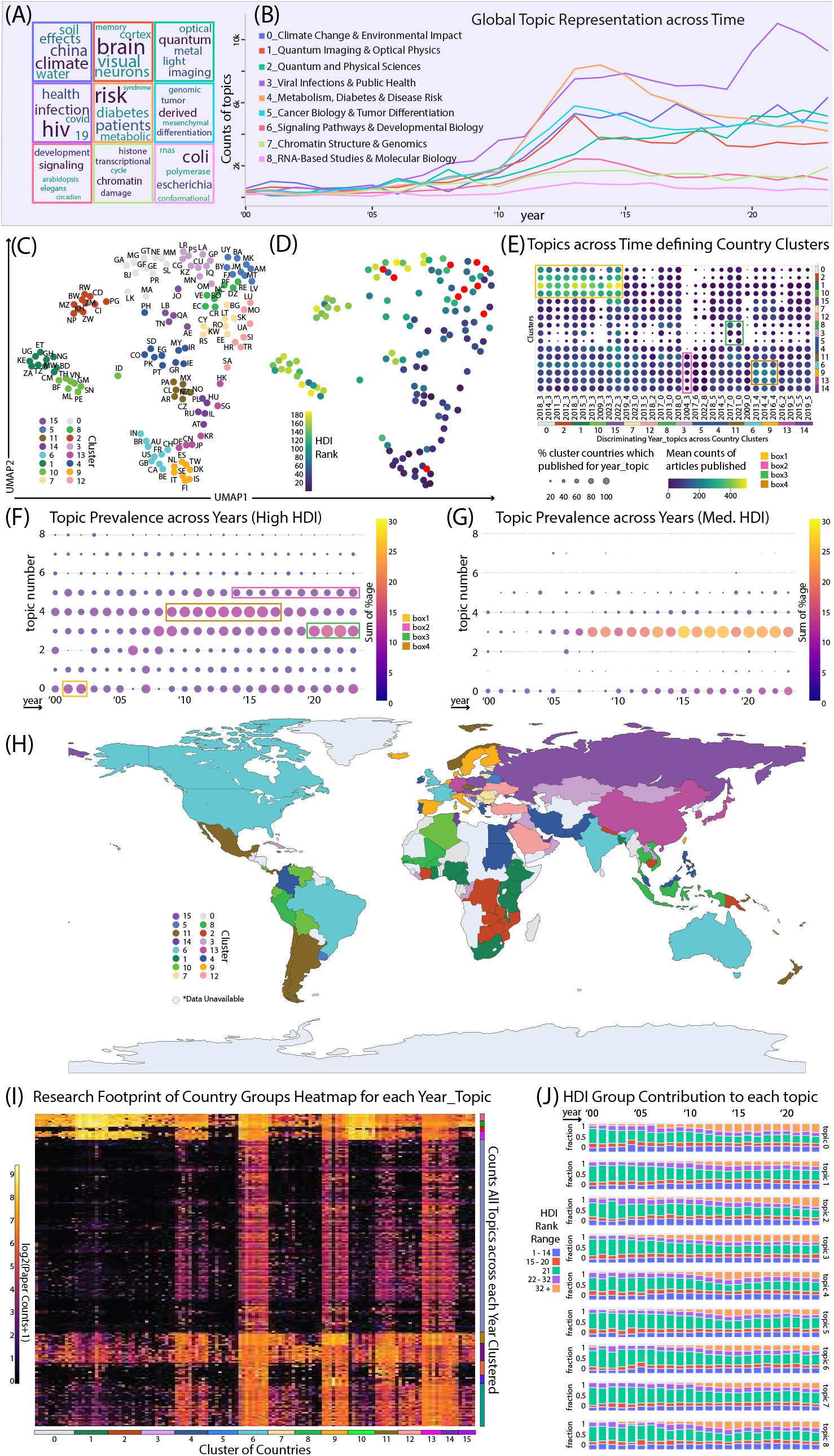
Topic Modeling of Scientific Research: Trends, Comparisons, and National Alignments. A – Key topics uncovered from papers in the life sciences co-authorship networks. B – Temporal analysis of these topics across all countries over time. C – UMAP and defined Leiden clusters of nations having similar research profiles. D – Clustering of countries by research trends and HDI alignment, highlighting the link between socioeconomic development and scientific priorities. E – Distinct research priorities and their evolution across country groups, highlighting the discriminative research focus of each cluster. F – Topic specific quantification of scientific contribution in high HDI countries across 2000-2023 G – Topic specific quantification of scientific contribution in medium HDI countries across 2000-2023. H – Holistic view of global scholarly research collaboration across the country clusters. I – Identifying influential country clusters guiding global research across multiple domains. J – Determining temporal shifts in research focus across countries in different HDI categories.

While overall scientific research output has grown exponentially over time (Suppl. Fig. 1), we investigated how this trend applies across disciplines (Fig. 3B, Supplementary Fig. 3B). We categorized each article into its most relevant topic. Unexpectedly, not all research areas show exponential growth, revealing striking disparities in scientific outputs among different topics. For example, Viral Infections & Public Health (Topic 3, Fig. 3B) surged post-2019 in response to COVID-19, whereas Metabolism, Diabetes & Disease Risk (Topic 4, Fig. 3B) declined after 2014. Following years (2000-2012) of concern and attention, Climate Change & Environmental Impact (Topic 0, Fig. 3B) plateaued over the last decade, likely due to shifts in funding policies. Meanwhile, Signaling & Developmental Biology (Topic 6, Fig. 3B), Chromatin Structure & Genomics (Topic 7, Fig. 3B), and RNA-Based Studies (Topic 8, Fig. 3B) have remained approximately stable across 24 years. An important observation is that the pandemic spurred significant growth in topic 3 (Viral Infections and Public Health), even as progress in other fields remained flat or declined over time.

Overall, we find nine major research areas prevalent globally over the past 24 years and capture shifting trends in scientific output. Our method offers a systematic and clear framework for monitoring the changing research landscapes, delivering important insights into the dynamics of scientific collaboration and advancement, which can assist in shaping research policies.

### Research interests align among nations with similar levels of human development

Leveraging the interpretable aspects of our approach, we mapped and inspect global research by linking articles to countries via author affiliations, revealing patterns in output and collaboration. Not only such insights can illuminate international partnerships but also highlight gaps in research areas, ensuring effective allocation of research and development resources.

Each country’s research repertoire was represented by its topic-wise paper count over 24 years. Next, we applied dimensionality reduction (Supplementary Figs. 3C-E) and defined Leiden clusters (Fig. 3C) of nations having similar research profiles. On scrutinizing each cluster, akin to cell annotation in single cell RNA-seq analysis, we observe that each cluster comprised of countries with comparable Human Development Index Rank (Fig. 3D). Strikingly, with no prior HDI information, our unsupervised approach de novo clustered nations with comparable HDI solely based on research trends. Of note, HDI encapsulates key factors like health, education, and living standards - all fundamental to research infrastructure. Our findings suggest that countries with similar HDI naturally converge on analogous scientific inquiries reinforcing the link between socioeconomic development and research priorities. Notably, while the UN does not provide HDI for all IBAN-defined regions (Supplementary Table 3), our approach enables HDI approximation based on research profiles. We posit that Taiwan’s development index aligns closely with Cluster 9 countries like Italy, the Netherlands, and Denmark, while Bosnia and Herzegovina share similarities with Cluster 5.

Drawing from the idea of analyzing specific cluster differences in single cell analysis, we identify primary research areas that distinguish each country cluster from the rest. This not only enhances the granularity of interpretability but also reveals key relationships between nations and their scientific priorities. Groups 6, 9, and 13 prioritized metabolic research (2013–2016, box4, Fig. 3E), while clusters 0, 1, 2, 10, and 15 were leading contributors to viral research, which has since declined (box1, Fig. 3E). Group 11 exhibited strong output in environmental studies (2017–2021, box3, Fig. 3E) but had minimal infectious disease research in 2004 (box2, Fig. 3E), setting it apart from group 3. Remarkably, high-HDI country groups (6, 9, 11, 13, 14) pursued broader scholarly agendas, likely enabled by greater resources and infrastructure. To further corroborate the hypothesis that research interests align with human development, we separate countries into high and medium HDI and quantified per topic scientific contributions over time as percentages. Our analysis reaffirms the striking trend: high-HDI nations showed a dynamic response to global challenges, shifting focus from climate change (2001–2002, box1, Fig. 3F) to diabetes (2009– 2018, box4, Fig. 3F) and then to viral infections post-2019 (box3, Fig. 3F). Their sustained interest in genomics after 2014, likely driven by Human Genome Project and CRISPR’s impact (box2, Fig. 3F), reflects their proactive approach to health innovation. In contrast, medium-HDI countries maintained a more stable focus on infectious and metabolic diseases (Fig. 3G).

Overall, these findings not only highlight global research gaps but also provides actionable insights for effective allocation of research expenditures. We conclude that a country’s scientific interests reflect its level of human development. Particularly, countries with medium HDI could greatly benefit from pooling resources and collaborating to address scientific challenges. In conclusion, our method lays the groundwork for crafting tailored scientific policies for each country and presents a thorough analysis of dynamic research patterns.

### Global collaborative patterns mirror the structure of genomic regulatory networks

Our scalable and interpretable approach offers an unprecedented overview of global scientific data, made possible by advances in organized data and computational capabilities. This allows us to explore patterns and trends that were previously difficult to detect. While one might expect neighboring regions to collaborate more closely, our analysis reveals that geographic proximity does not always align with shared scholarly interests. For example, while some African regions (Clusters 1 and 2, Fig. 3H) show a correlation between geographic proximity and related scientific topics, European countries demonstrate diverse research foci (Clusters 4, 6, 7, 9, 12; Fig. 3H). By analyzing paper counts over time, we identify key country clusters (Clusters 6, 9, 13, 14, Fig. 3I) that act like “lineage-defining transcription factors”—master regulators that dictate cell fate by activating specific gene programs and silencing others. Similarly, these nations shape the trajectory of global research, revealing where innovation is headed. This striking parallel between scientific collaboration and gene regulation offers a powerful, data-driven lens to gauge the future of research. Notably, COVID-related research has emerged as a key area of collective scientific effort. Thus, by demonstrating that geographic proximity does not always imply similarity in research topics, we highlight the complex and unpredictable dynamics driving international collaboration. This reinforces our approach to mapping the global scientific landscape based on topic modeling by each country (Fig. 3I).

Lastly, we categorized countries into five HDI groups (1-14, 15-20, 21, 22-32, 32+) to analyze how research focus shifts over time (Fig. 3J). Our findings show that high HDI countries (ranks 1-20) maintain stable research interests, indicating sustained agendas supported by strong resources. The U.S. (rank 21) demonstrates more dynamic shifts, reflecting its adaptability to changing scientific priorities. Countries ranked 32+ (China being a significant contributor) showed a significant rise in research output across topics post-2010, signaling a strategic shift and growing global scientific influence. In conclusion, our analysis provides a comprehensive understanding of the evolving research landscape, offering a robust structure to map the dynamics of scientific progress across different economic and geographical contexts.

### Global partnerships on the rise: the surge of cross-country collaborations

Research teams often form groups that have a similar scientific outlook, benefiting from shared resources, common contexts, and established networks. Moreover, as scientific endeavor become more interdisciplinary, researchers build larger communities to address these. However, while building these communities within the same country has some inherent advantages, cross-country collaborations often have the added benefit of orthogonal perspectives. Given this, we sought to test the relative trends of within country and cross-country collaborations over time. We categorized research articles as “within-country” if all authors were from the same nation and “cross-country” if multiple national affiliations were present. Juxtaposing cross-country and within-country publication counts, we find that indeed within-country research dominated until around 2005 (Fig. 4A, Supplementary Fig. 4A). However, this balance shifted notably post-2005, with cross-country collaborations accelerating sharply from 2010 and reaching significantly higher levels by 2023. To normalize for the number of publications, we also examined the proportion of cross-country research for each country (Fig. 4B, Supplementary Figs. 4B-D) and compared the relative outputs (as quantified by the area under the curve) for each year. Even after normalizing for the total number of publications, the same trend remains – cross-country collaborations are increasing over time. This underscores a fundamental shift in how biomedical research is conducted, reinforcing that scientific inquiry transcends geographical boundaries more than ever before.

**Figure 4.**
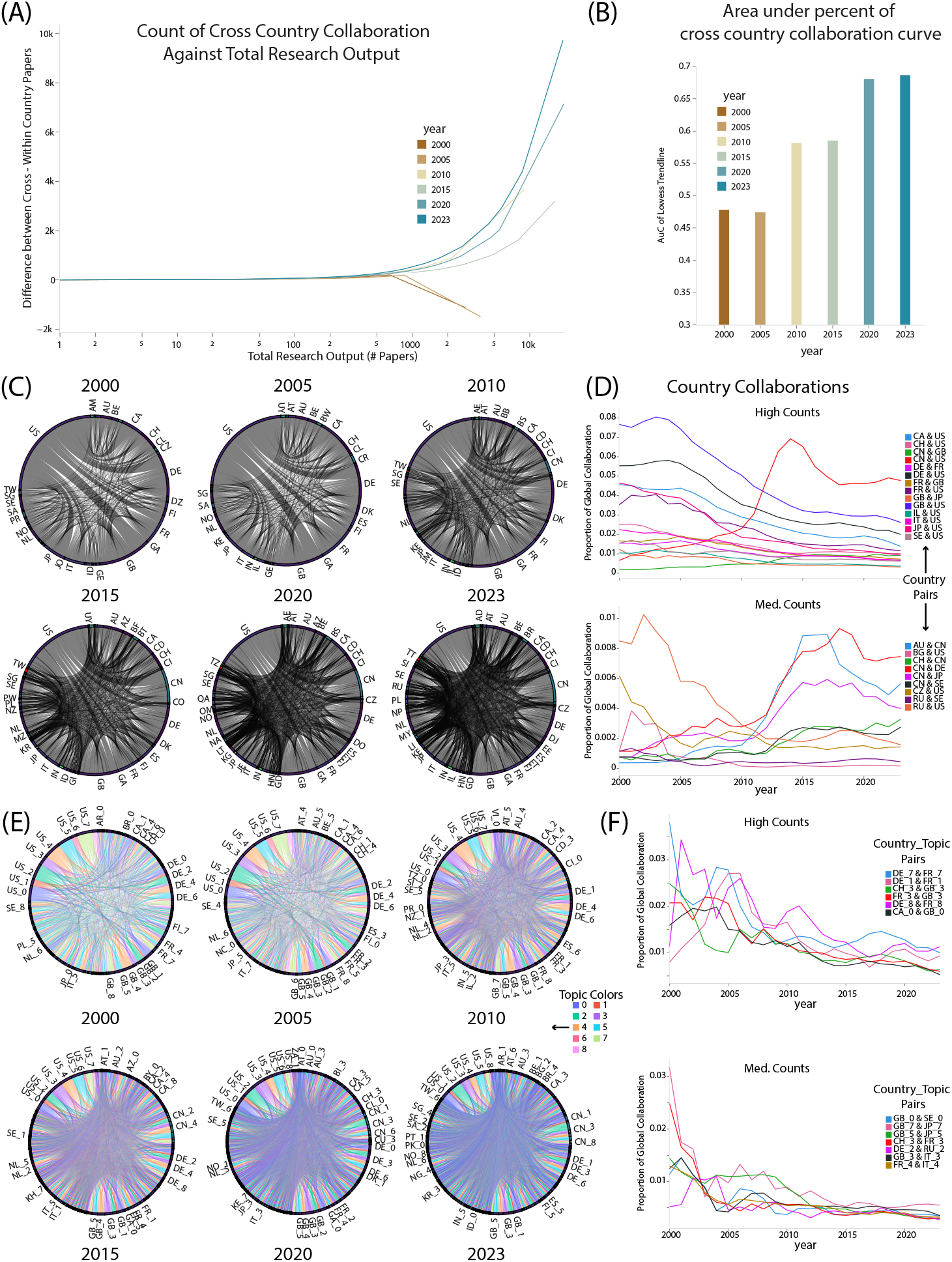
Shifts in the collaborative nature of global scholarly research over time: increasing global representation, democratization of research, and the growing need for strategic collaborations. A – Cross-country collaboration publication count vs. total research publications in 2000, 2005, 2010, 2015, 2020, and 2023. B – Area under the curve of cross-country collaboration percentage in 2000, 2005, 2010, 2015, 2020, and 2023. C – Contact maps of scientific collaboration of countries in 2000, 2005, 2010, 2015, 2020, and 2023. D – Scientific collaboration of countries across 2000-2023 in two groups based on the counts of collaborative papers (high and medium count country collaborations). E – Topic-based Contact maps of scientific collaboration of countries with high coefficient of variation in years 2000, 2005, 2010, 2015, 2020, and 2023. F – Topic-based scientific collaboration of countries with high coefficient of variation across years 2000 to 2023 in two groups based on the counts of collaborative papers (high and medium count country collaborations).

### Contact maps unveil emerging players in global scientific collaborations

After establishing the growing interconnectedness of scientific research, we next delved into country-specific patterns driving these partnerships. With more nations contributing to global research, we expected broader international engagement. Borrowing from genomic biology’s Hi-C contact maps—graphical representations of chromatin interactions within the genome derived from Hi-C sequencing to depict the frequency of physical contacts between different genomic loci—we defined scientific ties as any paper co-authored by researchers from two different countries. To improve resolution, we stratified these ties by publication counts.

Quantitatively analyzing shifts over time using coefficient of variation (CV), which here measures the variability of scientific output counts between countries, reveals a steep decline in the U.S.’s scientific partnerships with European nations such as Great Britain, France, and Sweden (Figs. 4 C-D). Surprisingly, despite geopolitical tensions, U.S.-China research ties have strengthened, whereas U.S.-Russia connections have weakened, likely at least partly driven by economic sanctions. Meanwhile, China is actively forging new alliances, shifting the epicenter of scientific innovation toward Asia. For instance, Switzerland’s research ties with the U.S. are declining, while its collaboration with China is expanding, alongside China’s increasing engagement with Denmark, Japan, Australia, and Sweden. Similarly, Great Britain’s once-strong connections with the U.S. and Japan have weakened.

While international scientific cooperation continues to grow, this shift is not uniform. Traditional leaders like the U.S. and Great Britain are increasingly being matched (and sometime outcompeted) by emerging powerhouses such as China, Japan, and Australia (Figs. 4C-D). The recent challenges to scientific funding in the U.S. may have more significant and far-reaching consequences, not just for U.S.-led biomedical research, but also for the global scientific community, given the interconnected nature of modern research. The rise of Asia as a central hub of scientific innovation represents a redistribution of intellectual capital, which could reshape funding priorities, policy decisions, and biomedical advancements. This shift also carries implications for industrial opportunities, with R&D-based companies seeking proximity to these emerging research powerhouses. Thus, understanding these evolving dynamics is essential for not just biomedical research but also economic growth in general.

### Democratization of research drives the need for strategic and topic-specific alliances

Given that our previous results (Figs. 4C-D) demonstrate clear shifts in country-specific collaborations, we sought to delve into this further at the level of specific scientific topics. Aligning partnerships with these priorities can enhance scientific impact and optimize resource allocation. Since specific research areas are prioritized by different countries (Figs. 3E-G), aligning partnerships with these priorities can enhance scientific impact and optimize resource allocation.

Consistent with the growing U.S.-China alliance in research, this trend spans multiple research domains (topics 2, 5, 6, 7; Supplementary Fig. 4E). However, across other research fields, nearly all other major U.S. partnerships with other countries have remained the same or gone down. This signals a systemic shift rather than a domain-specific change, highlighting a notable reduction in the U.S.’s collaborative research output (Supplementary Fig. 4E). When we examined other national partnerships, we observed that those with high coefficient of variation (CV) values were predominantly characterized by declining trends over time. This means that for many countries, the variability in their scientific collaborations was high, but these fluctuations were mainly negative, showing a downward trajectory. This decline in partnerships with high CV values suggests that, historically, research collaborations were more concentrated and stable, particularly among European nations (Fig. 4F). The sharp variability observed indicates that these nations had consistent and more focused collaborations in the past, but over time, these relationships became less stable and diminished. For example, partnerships such as Denmark– France (Topic 1, 7, 8) and Great Britain’s collaborations with France, Switzerland, and Italy in infectious disease research (Topic 3, Fig. 4F) have weakened. Notably, while global infectious disease publications have surged (Fig. 3B), inter-European research efforts in this field have declined. Similarly, Great Britain and Japan have seen a drop in joint genomics and signaling research output (Topic 7, 5; Fig. 4F). Lastly, a domain-specific analysis highlights the U.S.’s historical dominance in genomics (Topic 7), particularly after the Human Genome Project catalyzed global biomedical research. International collaborations were further driven by its large-scale initiatives like the ENCODE consortium which laid the foundation for global genomics research. However, the shifting dynamics and the challenges faced by science in US may start affecting future scientific breakthroughs. Ultimately, countries can forge impactful partnerships by focusing on prominent research areas and collaborating with leading contributors in those fields. Our findings highlight these evolving trends, offering a roadmap for nations to lower entry barriers and accelerate progress in emerging domains.

## Discussion

At a time when scientific research faces challenges from shifting U.S. governmental policies that have key implications for its foundation as a leader in funding education, training, and innovation, understanding the resilience and adaptability of global research networks is more critical than ever. The consequences extend beyond academia – undermining advancements in health, engineering, and technology. The implications also extend beyond countries as an attack on science transcends geographical boundaries.

However, the scientific community has demonstrated a capacity for self-sustenance, with life sciences research alone producing nearly two million publications in the past two decades and fostering unprecedented international collaboration. The scale of such data and the intricate nature of co-authorship networks posed significant challenges, necessitating the development of novel tools and methodologies to ensure both accuracy and interpretability in our findings, which can be addressed through interdisciplinary science—leveraging modern artificial intelligence techniques like natural language processing (NLP) in computer science and analytical methods for genomic data in computational biology and genomics to develop robust pipelines and analytical frameworks. By leveraging these interdisciplinary advancements, we reveal the shifting structure of global research communities and highlight the need to protect and strengthen these networks in the face of evolving geopolitical and funding landscapes.

Existing methods fall short in capturing the nuanced dynamics within these networks, primarily relying on topological features that provided limited insight into the deeper, underlying patterns of collaboration. We address these limitations by integrating advanced machine learning techniques, such as GATs^11^, BERT-based embeddings, zero-shot learning, and topic modeling, with scalable computational frameworks, as well as methodologies from genomic data analysis. By leveraging BERT embeddings, we were able to capture complex semantic relationships within paper titles, enhancing the model’s ability to understand and predict collaboration patterns. Additionally, our incorporation of zero-shot learning and TF-IDF provide alternative perspectives, enabling us to systematically compare more complex models like GATs versus simpler, more interpretable methods. This comprehensive approach allowed us to handle vast datasets efficiently while ensuring that the results remained both interpretable and meaningful, offering deeper insights into the evolving landscape of scientific collaboration.

One of the key strengths of this study lies in its ability to implement cluster-specific topic modeling, which captures information with a level of detail and clarity that surpasses traditional black-box methods. By focusing on specific clusters within the co-authorship networks, our approach allowed for a more granular analysis of the research topics and collaboration patterns that have emerged over time. This not only provided a clearer understanding of the evolution of scientific collaboration but also highlighted the critical role of high-quality embeddings in achieving robust predictive performance. Our analysis revealed several important trends, such as the significant peak in COVID-19-related research post-2020. These findings underscore the relevance of our methodology in mapping the evolution of global research priorities, offering a comprehensive view of how scientific focus shifts in response to global challenges and emerging technologies.

Furthermore, by linking topic-specific publications to authors’ affiliations over two decades and applying dimensionality reduction with Leiden clustering, we found that countries naturally grouped by research profiles align closely with their HDI ranks. Remarkably, our model de novo clustered nations with similar HDI solely based on research trends, reinforcing the link between scientific priorities and socioeconomic factors such as health, education, and living standards. Notably, our approach provides a data-driven estimate of HDI based on research activity, addressing gaps where the UN does not report HDI for certain IBAN-defined regions. Once again, data analysis methods from genomics played a crucial role, enabling us to analyze specific clusters and identify key research topics that differentiate country groups. Moreover, stratifying nations by HDI ranks revealed that high-HDI countries demonstrated a significant response to global challenges, actively contributing to health innovation, while medium-HDI countries maintained a more consistent focus on infectious and metabolic diseases. Also, our findings reveal that scientific collaboration is not solely determined by geographic proximity, as seen in the diverse research foci within countries despite their geographical closeness. Additionally, the identification of influential country clusters, particularly those resembling “master transcription factors,” underscores the pivotal role certain nations play in guiding global scientific trends. Finally, our analysis of HDI groups highlights the distinct shifts in research focus over time, with high HDI countries (ranks 1-20) maintaining stable agendas, likely due to stable resources and well-established scientific infrastructures. In contrast, the U.S. shows more dynamic shifts in focus, reflecting its ability to quickly adapt to emerging global challenges. Interestingly, emerging economies like China show a significant increase in research output post-2010, signaling a shift in global scientific influence, indicating a strategic shift and growing investment in specific research domains. This difference in research focus and adaptability likely stems from varying levels of resources, infrastructure, and perhaps differing national priorities. Through this meticulous analysis, we offer a powerful tool for interpreting co-authorship networks, revealing how scientific collaboration and thematic focus have evolved in response to global events and emerging challenges.

The shifts in research trends prompted us to examine how the nature of scholarly publications is evolving globally. By categorizing publications as “within-country” or “cross-country,” we found that countries are increasingly collaborating, even though within-country research still benefits from shared resources and established networks. This highlights that groundbreaking science, capable of global impact, requires international cooperation and strong research communities. Our analysis revealed several key partnership trends: a sharp decline in U.S. collaboration with European nations like Great Britain, France, and Sweden, while U.S.-China research ties have strengthened despite geopolitical tensions. Conversely, U.S.-Russia collaborations have diminished, likely due to economic sanctions. Notably, the U.S. is witnessing a broader decline in collaborative partnerships across research fields, except for its growing alliance with China, signaling a systemic shift in global scientific dynamics. This points to a broader trend where old scientific powerhouses are losing influence, and new collaborations are emerging, reducing the dependency on any single nation to drive global innovation. This is occurring at a time when cutting funding could have severe international consequences, particularly jeopardizing U.S. scientific leadership, especially as China rapidly increases its research funding and infrastructure.

Analyzing the intricate dynamics of global scientific collaborations and co-authorship networks from 2000 to 2023 presented significant challenges, requiring innovative tools and methodologies to ensure both accuracy and interpretability. To address this, we developed the “DISCo-Net” package, a powerful and flexible tool designed to seamlessly handle every stage of the analysis pipeline—from data embedding to model execution and visualization. The creation of DISCo-Net represents a major advancement in the analysis of large-scale scientific collaboration networks, enabling more nuanced insights into the evolving landscape of global research.

## Methods

### Introducing DISCo-Net, an open-source python package for scalable network curation and analysis

DISCo-Net is implemented in Python (version 3.10) and is designed to be used on compute clusters. While we provide extensive tutorials and documentation for Slurm workload manager-based Linux compute cluster, DISCo-Net is readily extendible to any other cluster configurations which are supported by dask^13^. DISCo-Net code is open source (under MIT License) and available to be installed via pip from Python Package Index (https://pypi.org/project/pydisconet/). Additional details, Jupyter notebooks and tutorials along with detailed descriptions are available on our GitHub page (https://github.com/jishnu-lab/DISCoNet).

### Parsing and constructing datasets using DISCo-Net

DISCo-Net currently curates and analyzes two datasets: Life Sciences fetched from OpenAlex^14^ data collection and Physical Sciences derived from arXiv research archives. The Life Sciences dataset is created using OpenAlex, an open source repository comprising 248 million documents derived from various sources including Microsoft Academic Graphs (MAG)^13^ and Crossref^16^. To avoid duplication of authors, OpenAlex implements its custom Author Name Disambiguation (AND) algorithm to uniquely assign author’s name, the institutions, paper concepts, citations attached to a work, the coauthors on a work, and an ORCID if available.

DISCo-Net performs Structured Query Language (SQL) queries via duckDB^17^ python package to fetch dataset from the OpenAlex’s Amazon Web Services (AWS) bucket, providing extensive scalability. Our approach allows us to perform server-side queries instead of fetching data via REST API which is typically capped at 100,000 requests per user per day or downloading the whole snapshot data which is more than 300GB and changes monthly. Specifically, we query:

**SELECT** id, display_name, publication_year, publication_date, primary_location, open_access, indexed_in, institutions_distinct_count, authorships FROM read_json(‘{s3_path}’)

**WHERE** type = ‘article’ AND is_paratext = false AND is_retracted = false AND publication_year >=2000 AND publication_year <=2023 TO ‘{save_path}/openalex_raw_data/{save_name}.json’ (FORMAT JSON);

In the above query, we fetch only research articles which are not retracted and are not paratext, i.e., front cover, back cover, table of contents, etc. Approximately 30GB of data satisfying the above query is fetched which consists of research articles across various disciplines from year 2000 to 2023. Of these, we subset research articles published in 90 journals which focus on biomedical / interdisciplinary research (Supplementary Table 4). Article counts for the years where a certain journal of interest did not exist, was set to 0. Since we are focusing on research collaborations, we filter out all the single author entries and drop all the authors which had no country of affiliation.

Next to create physical sciences research dataset, we download all the research papers cataloged in arXiv from Kaggle^15^. Unlike OpenAlex, arXiv data is not standardized and requires extensive preprocessing including AND. DISCo-Net performs AND pillaring on last and first_middle name entries derived from authors_parsed entities from each paper. In DISCo-Net’s AND algorithm we define last authors names as unique keys followed by fuzzy word name matching and acronym-based name merging. Specifically, we first remove all the authors containing collaboration, consortium, team or numbers in the author names. Subsequently, we use thefuzz python package for name matching, combining authors having more than 90 percent similarity into one. Lastly, in case of records with the same last name but possible combinations of acronyms of first and middle names were also merged into the same author. DISCo-Net extensively uses dask based distributed computing paradigm to perform this preprocessing to provide scalability. All subsequent analyses are performed on the research articles filtered by the pre-processing.

### Creation of undirected weighted co-authorship networks for each dataset

For each year *y* in {*y* ∈ ℤ ∣ 2000 ≤ *y* ≤ 2023}, we create an undirected weighted co-authorship network in which each node of the network is an author ∈ {1, 2, …, *N*} where *N* is the total number of unique authors which published in year *y*. Mathematically, this network can be represented by its adjacency matrix *A*, where for each author pair *i, j, A*_*ij*_ = *A*_*ji*_ = 1 ∗ *c*_*ij*_ if they publish a research article together, else *A*_*ij*_ = *A*_*ji*_ = 0. Here, *c*_*ij*_ = *c*_*ji*_ ∈ ℕ represents the counts of papers published by author pair *i, j* together and defines the edge weight between the author pair. Lastly by construction, we don’t any self-loops, i.e., *A*_*ii*_ = 0.

Many real-world networks including co-authorship networks are known to be scale-free following the power law^16^ *P*(*K*) ∝ *K*^−γ^; where *P*(*K*) is the probability that a randomly chosen node has degree (i.e., the number of connections or edges a node has) *K. γ* > 0 is the power-law exponent typically varying from 2 to 3 for scale free networks. Upon fitting linear regression to the power law, log(*P*(*K*)) = −*γ* + *a* log(*K*) with *a* being the proportionality constant, we compute *R*^2^ across all the years. *R*^2^ measures the proportion of the variance in the dependent variable log(*P*(*K*)) that is predictable from the independent variable log(*K*) in the model and provides insights into the fit.

### Topological analysis of co-authorship networks

Various network topology characteristics for the built networks are calculated using NetworKit^17^ python package, which has approximation algorithm in in case where calculation of exact values is computationally unwieldy. Particularly, we calculate:

1. Network diameter which is maximum length of the shortest paths between any two nodes in the network.
2. Average degree centrality which measures the average connectivity of nodes in a network. It is given by 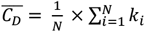, where *i* is an author in a year, *N* is the total number of authors and *K*_*i*_ represents degree of author *i*.
3. Average local clustering coefficient measures how close the node’s neighbors are to being a complete graph and is given by 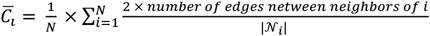. Here, 𝒩_*i*_ represents the set of neighbors of *i* and |. | is the cardinality operator.
4. Exact global clustering coefficient captures tendency of a network to form triangles.

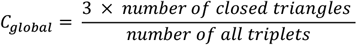

### Featurization of authors for each dataset and year

Every author in the co-authorship network is described by a feature list which is a concatenation of all the titles of papers published by the author in that specific year. Concretely, for each author *a*_*i*_ publishing *P*_*i*_ papers in a specific year *y*, we get a feature list 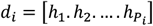 where *h* represents the heading or title of the specific paper. Subsequently, we generate a 768 dimensional feature vector for each author by two distinct approaches.

For the first one, we use a pretrained sentence BERT model trained on abstracts from PubMed and full-text articles from PubMedCentral followed by fine-tuning on Microsoft Machine Reading Comprehension (MS MARCO) dataset from Hugging Face^18–20^, which enables the model to have better semantic understanding in our context. Feature list for each author *i* – (*d*_*i*_) acts like a paragraph for sentence transformer with each title being a sentence. Each sentence is first tokenized by the model tokenizer and then each token is embedded to a 768 dimensional feature vector. By the virtue of how BERT based transformers are trained, each sentence’s token is prepended by [*CLS*] token which captures the overall context of all the tokens and hence the sentence itself. Hence, we use the embedding of this [*CLS*] token to describe each author. In the end, for a year *y* having *N* authors we get a feature vector matrix 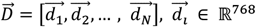 and 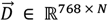.

In the second approach, we implement Term Frequency – Inverse Document Frequency (TF-IDF) based featurization of the author feature list *d*_*i*_. TF-IDF evaluates the importance of a word in a document relative to a collection of documents. While the TF-IDF value increases proportionally to the number of times a word appears in the document but is offset by the frequency of the word in the entire corpus. TF-IDF value for a term *t* in a document *d* within a corpus *D* is calculated as *TF* − *IDF*(*t, d*_*i*_, *D*) = *TF*(*t, d*_*i*_) × *IDF*(*t, D*). *TF*(*t, d*_*i*_) measures how frequently a term *t* occurs in a document *d*_*i*_ and is calculated by 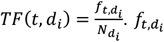 is the raw count of term *t* in document *d*_*i*_ and 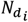 is the total number of terms in document *d*_*i*_. *IDF*(*t, D*) measures how important a term is in the entire corpus. It is calculated as 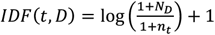, where *N*_*D*_ is total number of documents in the corpus *D* and *n*_*t*_ is the number of documents in which the term *t* appears.

First for a year *y*, we define the corpus by tokenizing titles of all the papers published (*D*) using a custom lemmatizer based on spaCy’s *en*_*core*_*web*_*sm* NLP^21^ model. To filter out noise during lemmatization, we only account for words having length greater than 2, convert all the nouns to their root form using TextBlob, and remove any punctuation or stop words. Following tokenization, we fit the TF-IDF model using scikit-learn’s pipeline comprising of CountVectorizer() and TfIdfTransformer(). Then for each author, we transform its feature list using the trained model to yield a feature vector matrix 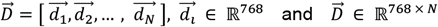 key hyperparameters of interest which we choose are:

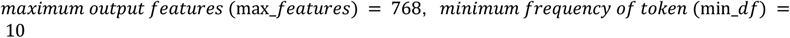

We choose the output dimension of vectors to be 768 to make it comparable to BERT based featurization. min_*df* avoids extremely rare words to be given very high importance, thus balancing frequency and rarity to describe each document *d*_*i*_.

### Defining the groups of authors based on productivity and type of collaborations

We stratify authors into four groups in two ways based on the number of papers published by them (*C*), node betweenness (*C*_*B*_) of the author and their degree in the network (*K*).

In the first approach, we classify authors based on their paper count (number of papers) and degree (frequency of collaboration). Number of papers published by each author within a specific year reflects productivity of authors in a specific year while degree reflects the number of collaborations of the author. We threshold the number of papers published at 98th percentile (*p*_1_) and degree at 95th percentile (*p*_2_). Therefore, we get 4 groups as:

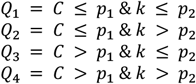

For the second grouping, we clump authors based on their nature/quality of alliances (within cliques or bridging groups) and degree. Nature of collaborations of an author is determined by their node betweenness which captures the importance of a node (here, author) by calculating the number of shortest paths passing through a node *i*, and is defined by:

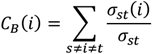

where, *σ*_*st*_ is the total number of shortest paths from node s to node t and *σ*_*st*_(*i*) is the number of those shortest paths that passes through node *i*. Again, we threshold the node betweenness at 98th percentile (*p*_1_) and degree at 95th percentile (*p*_2_) to get:

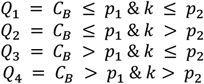

### Setting up link prediction task for ML methods

We use link prediction as our Machine Learning task. More formally, given a pair of authors and their feature vectors, we aim to predict if they collaborated or not, i.e. if an edge exists in the coauthorship graph. The network is split into train, validation and test sets in a 0.6 : 0.2 : 0.2 ratio, with no common edges to avoid data leakage. Performance metrics are reported for the whole set and for the individual groups defined based on productivity and connectivity. To ensure scalability, we sub sample the local neighborhood instead of loading the network into memory.

### Training graphical attention network on link prediction task

We train graph attention networks in edge prediction setting using PyTorch Geometric (PyG)^22^. Input to GAT model is author features derived from BERT based embedding 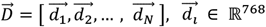 and 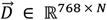 which are transformed using masked self-attention.

In summary, each node (author) feature is linearly transformed into a 256 − *dimensional* reduced embedding space parametrized by shared weight matrix, *W* ∈ ℝ^256 × 768^. This is followed by *LeaKyReLU* transformed self-attention score *e*_*ij*_, which are generated by a shared fully connected linear layer, *a* ∈ ℝ^256 × 256^ → ℝ. *e*_*ij*_ indicates the importance of authors *j*’s features to author *i* and is computed only for neighbors 𝒩_*i*_ of author *i*: *j* ∈ 𝒩_*i*_ in the co-authorship network. Final attention score *α*_*ij*_ is normalized across all possible neighbors *j* by using the *softmax* function:

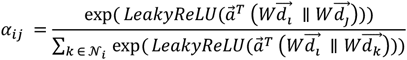

where *T* represents transposition and ∥ is the concatenation operation. New feature vector for author 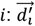 is then generated by linear combination of the neighborhood features corresponding to normalized attention scores:

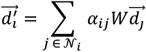

Our GAT model is trained for 4 epochs, has 2 attention layers with 1 attention head and outputs 256 − *dimensional* updated embeddings.

GAT model is trained with the objective of reconstructing the adjacency matrix and hence the initial network itself. For each author *i* in the network we subset local neighborhood using LinkNeighborLoader() function of PyG and take 2, 3 and 1 first-, second- and third-degree neighbors respectively to define a local graph. All the true edges are labelled 1 with the collaborating authors forming 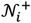 and equal number of negative edges are sampled and labelled 0 with non-collaborating authors forming set 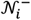. After updated embeddings are generated from GAT, we calculate the loss *L* as:

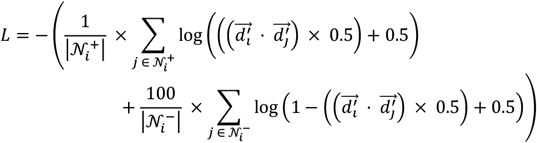

|. | represents cardinality of set. For optimization we use Adam^23^ optimizer with learning rate of 0.01 and weight decay of 0.001 and calculate performance of the model using Area under Receiver-operating characteristic curve (AuROC).

### Zeroshot modeling using embeddings of author features

In zeroshot modeling, we leverage already generated embeddings to perform link prediction for authors in the network. This is performed on embeddings generated by both BERT based method and TF-IDF. We do not require any training and hypothesize that the embedding’s themselves are rich enough to capture research and co-authorship context. Hence, for each author *i* in year 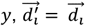. Utilizing the same loss function as in GAT modeling, we generate predictions on the sampled network of the author and calculate Area under Receiver-operating characteristic curve (AuC).

We turn to topic modelling frameworks to derive interpretable insights, particularly into research topic dynamics from a temporal perspective. We use the BERTopic framework which consists of three major steps.

Firstly, titles of research articles from each year *y* are embedded into a 768-dimensional space using a sentence transformer. The resulting feature matrix 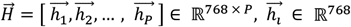 (where *P* is the number of papers published in year *y*), undergoes dimensionality reduction via Uniform Manifold Approximation and Projection (UMAP)^27^ to 32 dimensions. Topics are then extracted using Hierarchical Density-Based Spatial Clustering of Applications with Noise (HDBSCAN), which identifies stable clusters with robust outlier detection. HDBSCAN^28^ hyperparameters are optimized for domain specificity:

For Life Sciences:

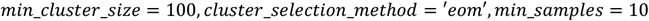

For Physical Sciences:

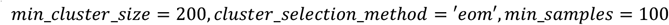

Secondly, To define topics, we implement class-based TF-IDF (c-TF-IDF) via BERTopic, treating all titles in a cluster as a single document. The term frequency is given by 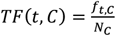 and inverse document frequency is adjusted as 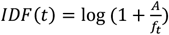 where *A* is the average word count per class. The c-TF-IDF score is then computed as:

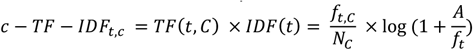

And lastly, Topics across years are merged if their *cosine similarity* ≥ 0.8, refined using a custom lemmatizer and CountVectorizer, and reassigned to documents. Each topic is represented by its top 50 terms, with word clouds generated from the five highest-weighted terms. Finally, dynamic topic modeling is performed by applying the trained c-TF-IDF model iteratively across yearly subsets, incorporating evolutionary tuning where each year’s topic distribution is averaged with that of the preceding year. This approach enables the robust temporal tracking of research themes.

### Construction of the Country-Topic Matrix

To analyze global research trends, we constructed a country-by-topic matrix based on the number of publications in each research topic per year from 2000 to 2024. We obtained country-wise topic distributions from the OpenAlex dataset and filtered out missing values and unassigned topics (−1). The dataset was structured such that each row represented a country, and each column represented a unique topic-year combination.

Formally, we defined the country-topic matrix *X* as:

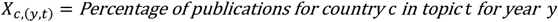

where:

*C* is the set of all countries,

*Y* = {2000, …, 2024} is the set of years,

*T* = {0,1, …, 9} is the set of topic labels (excluding −1),

The percentage is computed using log-scaled normalization:

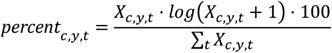

where *X*_*c,y,t*_ represents the raw count of publications in topic *t* for country *c* in year *y*. Countries with fewer than 10 total publications per year were excluded.

### Postprocessing of Country-Topic Matrix

The country-topic matrix was transformed into a high-dimensional feature space, where each row corresponded to a country and each column corresponded to a unique (year, topic) combination. The data was then standardized using Z-score normalization:

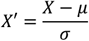

where *µ* and *σ* are the mean and standard deviation of each feature, respectively. This ensures that features are on a comparable scale before downstream analysis.

Next, we performed dimensionality reduction using Principal Component Analysis (PCA) to capture the most informative variance in the data:

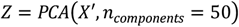

A k-nearest neighbor (kNN) graph was then computed based on the PCA-reduced space:

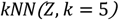

We then applied Leiden clustering; a community detection algorithm optimized for graph-based structures:

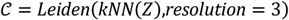

where 𝒞 represents the discovered clusters. A choropleth map was generated to visualize the spatial distribution of Leiden clusters. To visualize the clusters, we embedded the data into a two-dimensional UMAP space with an adjusted spread and min-distance for better separation:

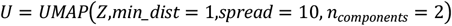

### Cluster Characterization via Differential Topic Analysis

To identify the distinguishing topics within each cluster, we performed differential expression analysis akin to single-cell RNA sequencing. Specifically, for each cluster *K*, we used the Wilcoxon rank-sum test to compute differentially enriched topics (DETs):

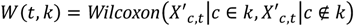

Topics were considered significantly enriched in a cluster if they met the following criteria:

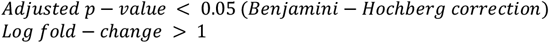

The top DETs per cluster were used to interpret the dominant research themes associated with different country groups.

Subsequently, to examine the temporal evolution of dominant research topics across countries with varying levels of Human Development Index (HDI), we generated bubble plots where the size of each bubble represents the number of countries for which a specific research topic was the most prevalent each year. This approach provides insights into how research priorities differ between high-HDI (Rank 1 to 50) and medium-HDI (Rank 100-150) nations over time. The dataset was first grouped by country, year, and research topic (work_name) to calculate the total number of publications in each category. The relative proportion of publications per topic within a given country-year pair was computed as follows:

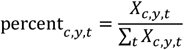

where *X*_*c,y,t*_ represents the number of publications in topic *t* for country *c* in year *y*. This normalization accounts for variations in total publication output across countries and years. To identify the most dominant research topic for each country each year, we determined the topic with the highest publication count within that country-year pair. Subsequently, we computed the number of countries in which each topic was dominant for every year. The resulting dataset provided a temporal distribution of leading research themes across different HDI groups.

### Hierarchical Clustering of feature space

To analyze the relationship between topics and countries, hierarchical clustering was applied to reorder a document-topic matrix. We used Ward’s method for column clustering:

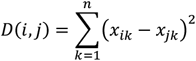

where *D*(*i, j*) represents the distance between two clusters *i* and *j*. Cluster assignments were computed using:

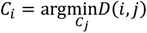

A heatmap was then generated, with Leiden clusters mapped to color annotations.

### Human Development Index (HDI) Categorization

Each author’s country was mapped to its HDI rank, and countries were grouped into bins based on predefined HDI thresholds. The percentage distribution of works per HDI category was computed as:

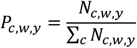

where *P*_*c,w,y*_ represents the proportion of works in category *c* for a given work *w* and year *y*, and *N*_*c,w,y*_ is the count of works in that category. This allowed the creation of a stacked bar chart showing the evolution of HDI-based work distribution over time.

### Analyzing Cross-Country Research Collaboration Trends

In our research, we analyze cross-country collaboration patterns over time by processing data from multiple years, specifically from 2000 to 2024, sourced from OpenAlex. For each year, we read the relevant paper titles and author information, then preprocess the data to focus on the collaboration details between countries. First, we filter and group the data by paper titles, creating a list of countries associated with each paper. Using this, we compute the number of collaborations between countries and within each country by applying the following two key measures:

1. Cross-country collaborations: This is calculated by counting the instances where a country is involved in a paper with authors from other countries. Formally, for a given country *c*, the number of cross-country collaborations, denoted as *C*_*c*_, is:

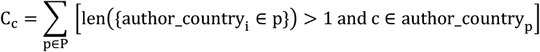

where *P* represents the set of all papers, and author_country_*p*_ is the set of countries for paper *p*.
2. Within-country collaborations: This counts papers with authors only from a single country *c*, denoted as *W*_*c*_, calculated as:

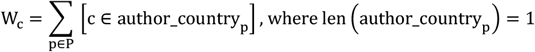

Using these counts, we compute the collaboration count difference *D*_*c*_ and collaboration percentage *P*_*c*_ for each country *c* in each year:

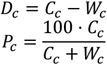

These metrics are then visualized with scatter plots, showing the relationship between the total research output (number of papers) and the percentage of cross-country collaborations, as well as the absolute difference in collaboration counts. A Lowess regression is applied to capture the trend in these relationships across time. Additionally, we compute the Area under the Curve (AuC) for each year to quantify the overall collaboration trend, where the AuC is given by:

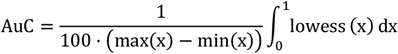

### Analyzing Pairwise Interactions Over Time using Coefficient of Variation

We utilized Circos plots to visualize cross-country research collaborations over time using data from OpenAlex for the years 2000 to 2023. The dataset was preprocessed to identify co-authorship relationships between countries by grouping authors based on their work and filtering for collaborations involving multiple countries. Each collaboration was then mapped to specific research topics using a topic modeling approach. For each year, we generated two types of Circos plots: one representing topic-specific collaborations and the other showing overall co-authorship trends. These plots were constructed using the pycirclize package, where the color and width of the links were customized to reflect topic-specific or general collaboration frequencies.

To analyze the pairwise interactions between research topics over time, we compute the normalized interaction count for each year and each pair of entities. The interaction count for each pair was normalized by dividing it by the total interaction count for that specific year to account for temporal variations in the data. This normalization process can be expressed mathematically as:

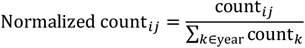

where count_*ij*_ represents the interaction count between entities *i* and *j* in a specific year, and the denominator is the total count for the year.

To ensure that the trends for each pair of entities across years are robust, a moving average (using a window of 3 years) was applied to smooth the interaction data:

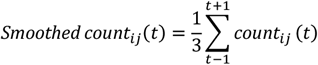

Next, we assessed the variability of the interactions using the Coefficient of Variation (CV), which measures the relative variability in relation to the mean. The CV is calculated as:

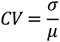

where *σ* is the standard deviation and *µ* is the mean of the smoothed interaction count. A lower CV indicates more stable interactions over time, while a higher CV suggests greater variability. Based on the CV, we applied thresholds to filter pairs of entities with low stability (high CV). Specifically, pairs with a CV less than 0.3 (or a mean value below 0.005) were considered stable enough to include in the final analysis for the “high mean counts” filter. For the “low mean counts” filter, pairs with a CV less than 0.4 or a mean above 0.01 or below 0.005 were excluded. The thresholds for these filters were adjusted to focus on more stable and significant interactions.

Once valid pairs were identified based on the CV and mean value thresholds, we plotted the interaction trends over time. The interaction data for each pair were visualized using line plots, with the x-axis representing years and the y-axis representing the smoothed interaction counts.

## Supporting information

Supplementary_Figure_Combined

## Code and Data Availability

DISCo-Net code is open source (under MIT License) and available to be installed via pip from Python Package Index (https://pypi.org/project/pydisconet/). Additional details, Jupyter notebooks and tutorials along with detailed descriptions are available on our GitHub page – https://github.com/jishnu-lab/DISCoNet.

## Supplementary Figure Legends

**Supplementary Figure Legends Supplementary Figure 1**

A – Number of published papers, number of authors and number of co-authorship links across 2 decades for manuscripts in the life sciences.

B – Number of published papers, number of authors and number of co-authorship links across 2 decades for manuscripts in the physical sciences.

**Supplementary Figure 2**

A – Schematics of network topological metrics – network diameter, average node betweenness, average degree centrality, and global clustering coefficient.

B – Network topological metrics (network diameter, average node betweenness, average degree centrality, and global clustering coefficient) of co-authorship networks from 2000-2023 in the life sciences.

C – Network topological metrics (network diameter, average node betweenness, average degree centrality, and global clustering coefficient) of co-authorship networks from 2000-2023 in the physical sciences.

D – AUCs of co-authorship link prediction by the GAT model from 2000-2023 in the physical sciences. Panels reflect performance overall, performance for different co-authorship patterns as defined by stratification using degree and node betweenness, performance for different co-authorship patterns as defined by stratification using degree and number of papers published.

E – AUCs of co-authorship link prediction by the BERT model from 2000-2023 in the physical sciences. Panels reflect performance overall, performance for different co-authorship patterns as defined by stratification using degree and node betweenness, performance for different co - authorship patterns as defined by stratification using degree and number of papers published.

F – AUCs of co-authorship link prediction by the TF-IDF model from 2000-2023 in the physical sciences. Panels reflect performance overall, performance for different co-authorship patterns as defined by stratification using degree and node betweenness, performance for different co-authorship patterns as defined by stratification using degree and number of papers published.

**Supplementary Figure 3**

A – Key topics uncovered from papers in the physical science co-authorship networks. B – Temporal analysis of these topics across all countries over time.

C – Principal components of dimensionality reduction of nations having similar research profiles. D – PCA plot of nations having similar research profiles.

E – Clustering of countries by research trends and HDI alignment

**Supplementary Figure 4**

A – Cross-country collaboration publication count vs. total research publications in 2000-2023.

B – Area under the curve of cross-country collaboration percentage from 2000 to 2023.

C – Cross-country collaboration percentage vs. total research output in 2000, 2005, 2010, 2015, 2020, and 2023.

D – Cross-country collaboration percentage vs. total research output from 2000 to 2023.

E – Topic-based scientific collaboration of the U.S. with high count country from 2000 to 2023.

F – Topic-based scientific collaboration of the U.S. with medium count country from 2000 to 2023.

**Supplementary Table 1**

Words and corresponding topics for Biomedical Sciences (OpenAlex) dataset

**Supplementary Table 2**

Words and corresponding topics for Physical Sciences (Arxiv) dataset

**Supplementary Table 3**

HDI ranks and associated countries as inferred from UN

**Supplementary Table 4**

List of curated journals for which scientific research articles were fetched and analyzed

