## Supplementary figures and images for "Cracking the code of co-authorship networks geo-temporally using interpretable machine learning"

### Supplementary_Figure_Combined

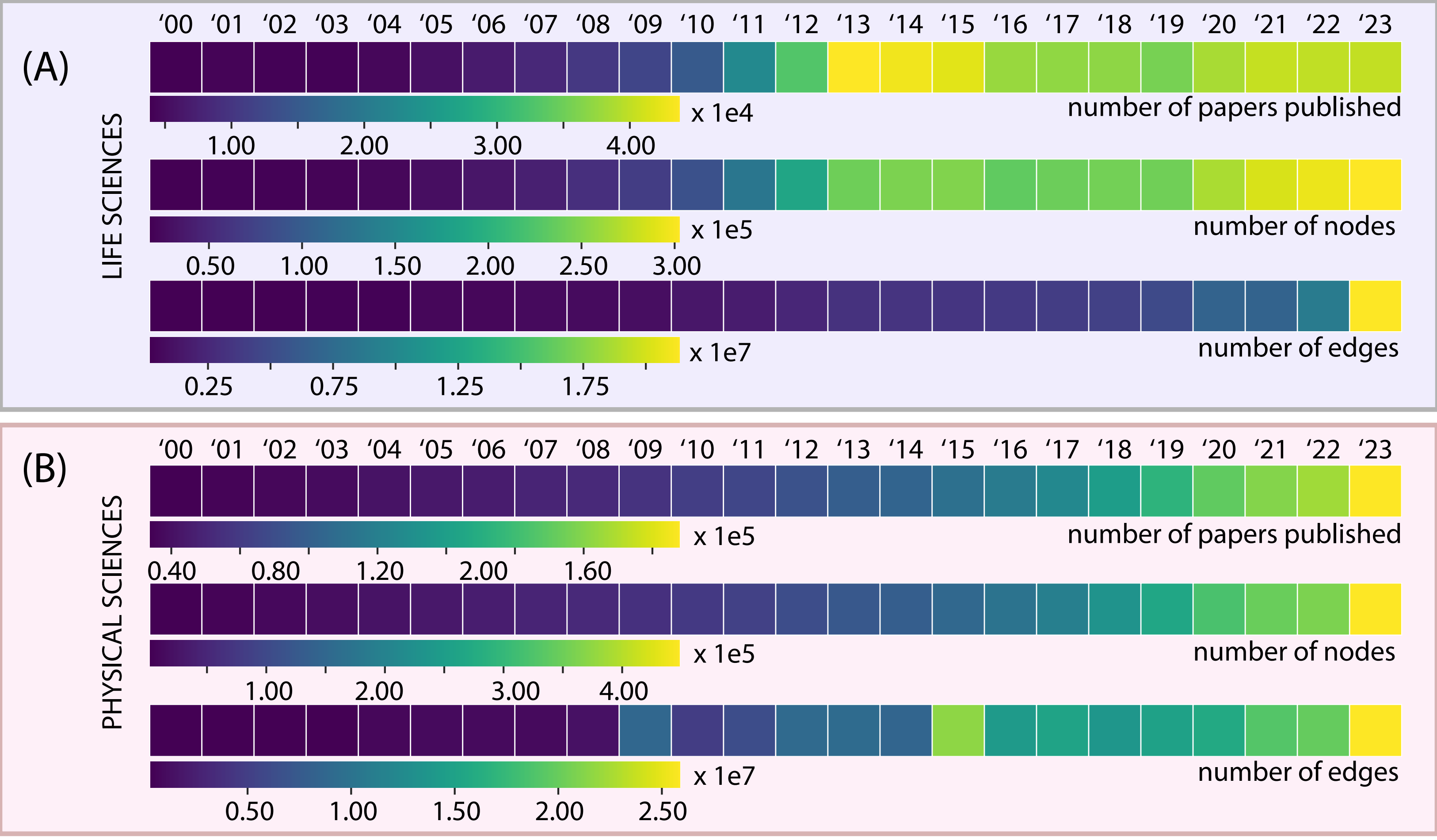

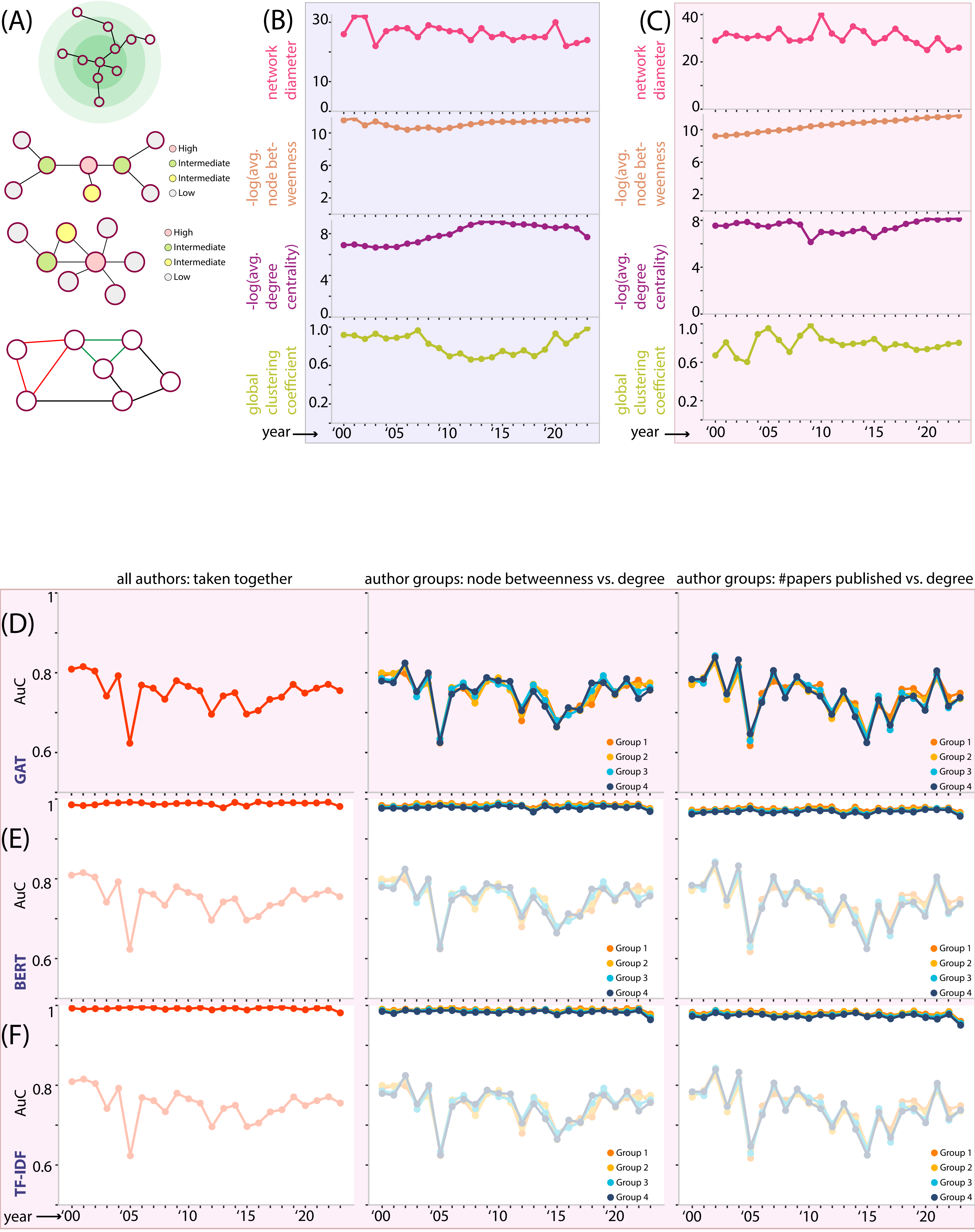

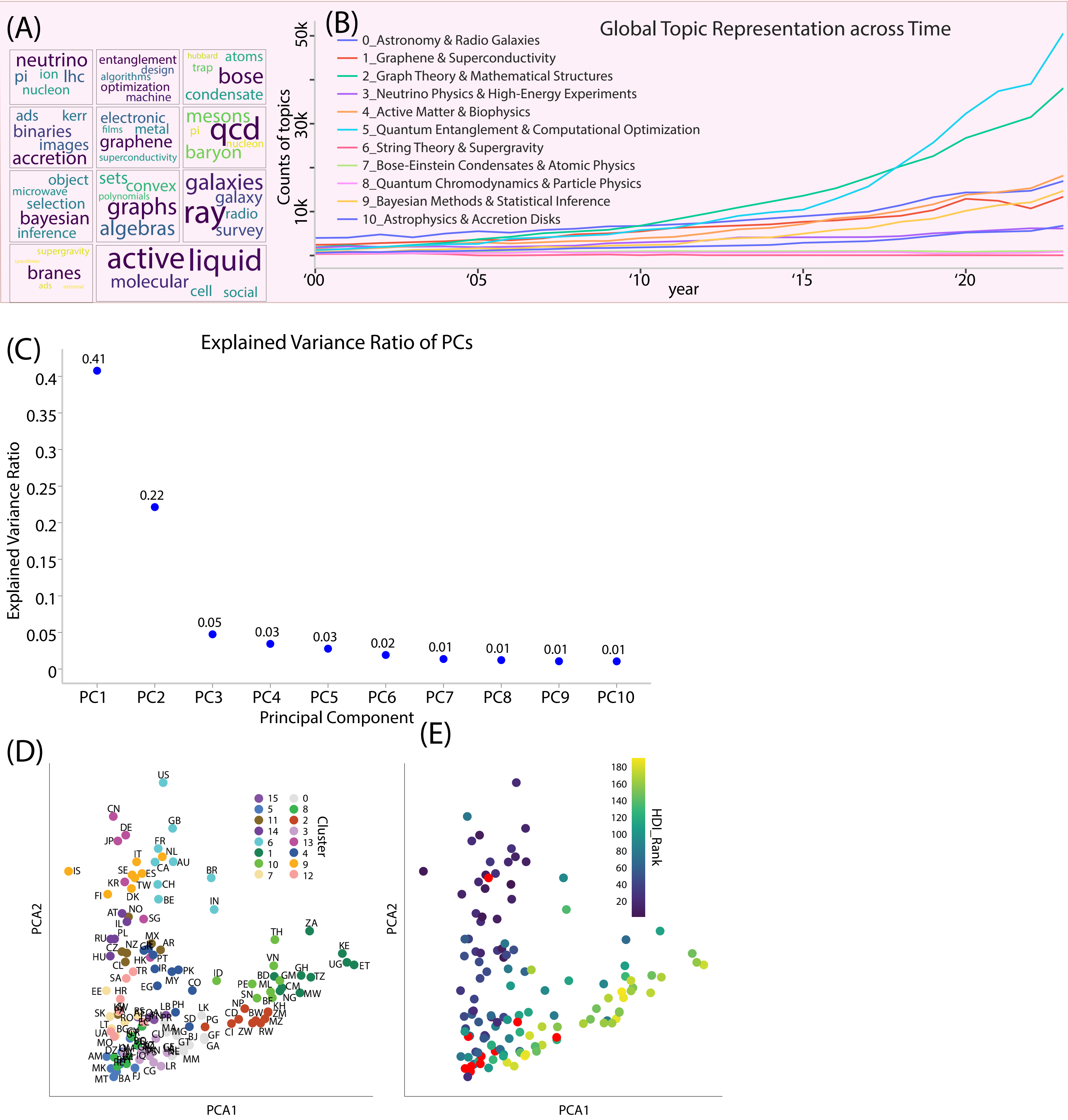

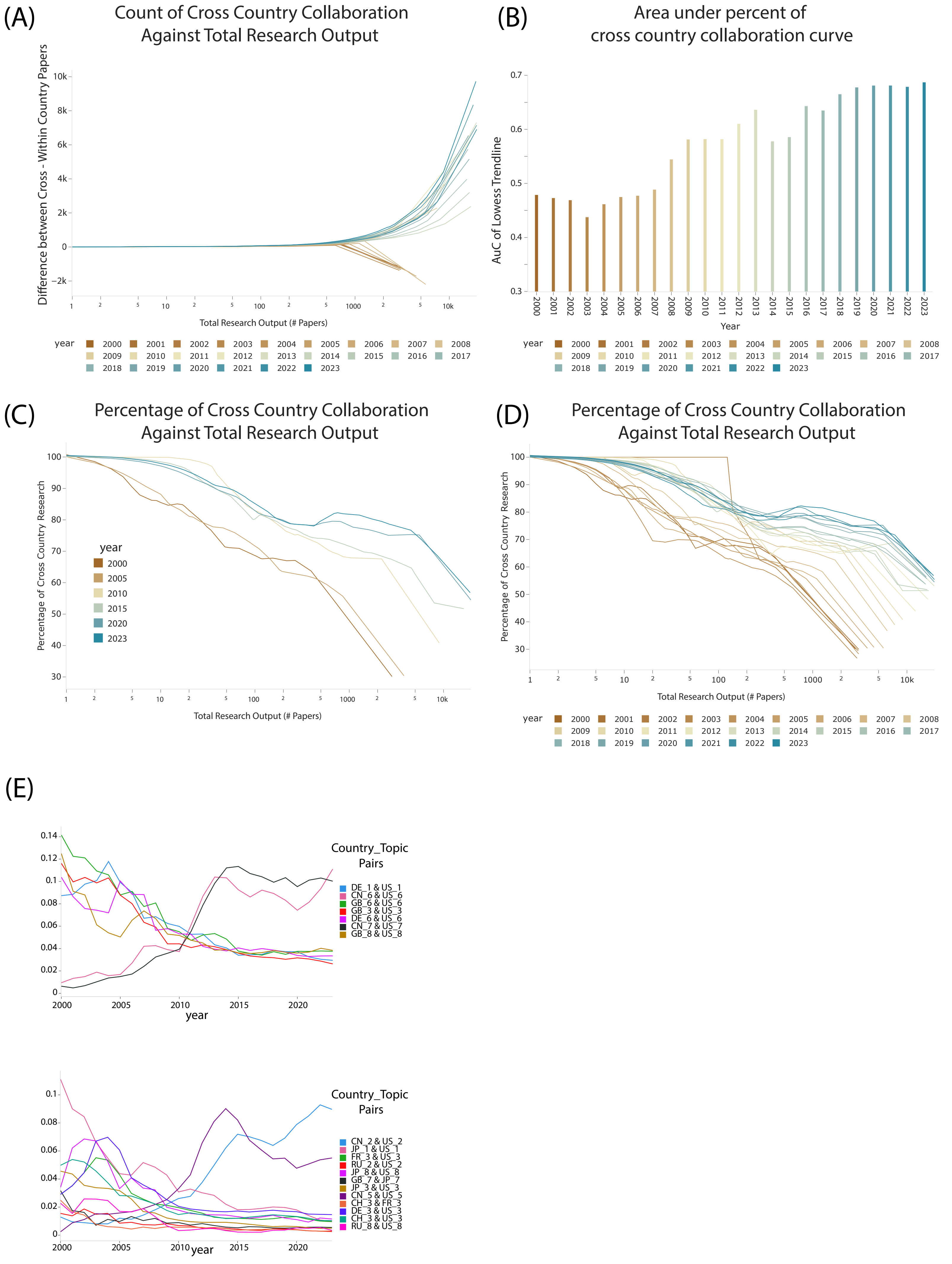

### Supplementary_Video_1

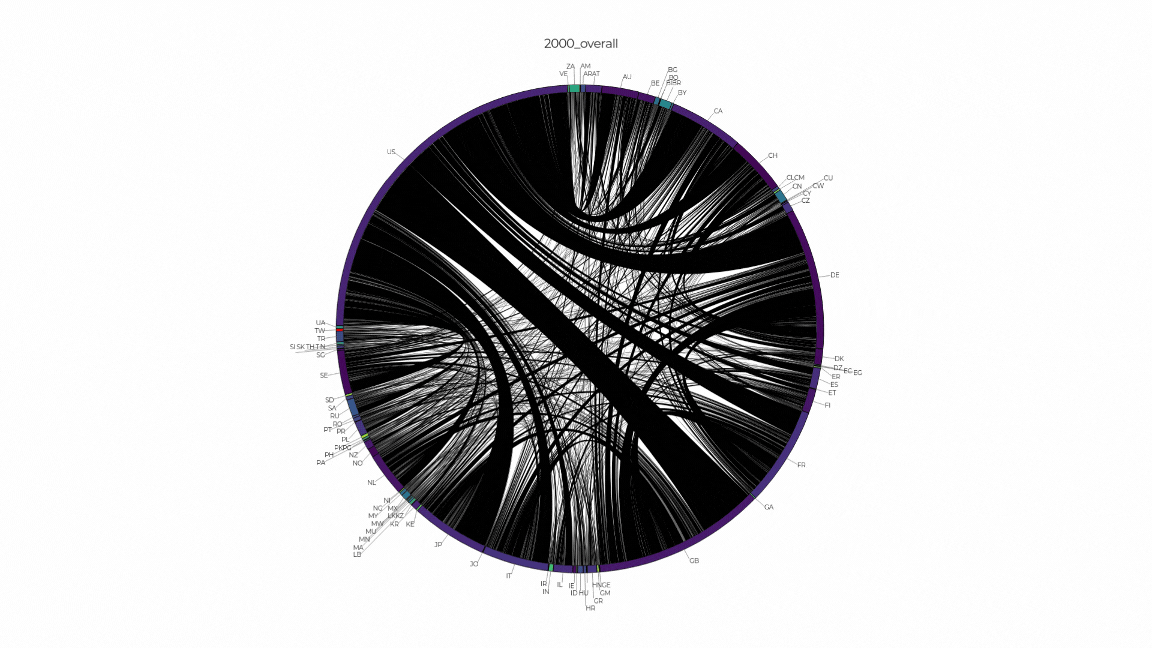

### Supplementary_Video_2

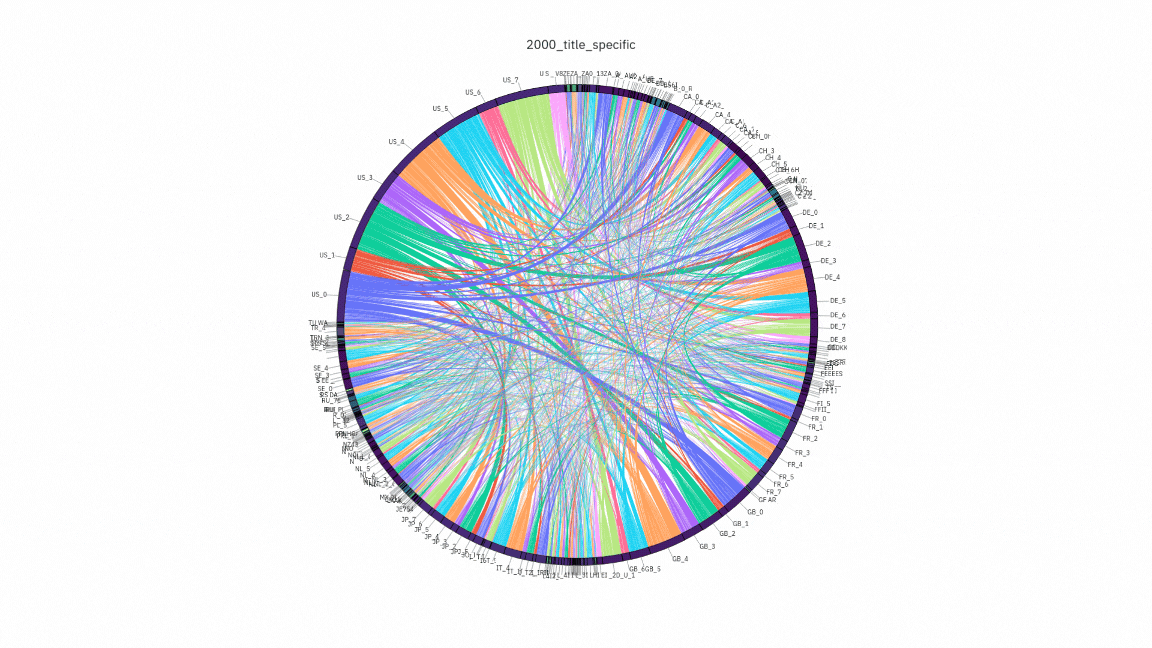
